# Feeding Rates in Sessile versus Motile Ciliates are Hydrodynamically Equivalent

**DOI:** 10.1101/2024.05.15.593824

**Authors:** Jingyi Liu, Yi Man, John H. Costello, Eva Kanso

## Abstract

Motility endows microorganisms with the ability to swim to nutrient-rich environments, but many species are sessile. Existing hydrodynamic arguments in support of either strategy, to swim or to attach and generate feeding currents, are often built on a limited set of experimental or modeling assumptions. Here, to assess the hydrodynamics of these “swim” or “stay” strategies, we propose a comprehensive methodology that combines mechanistic modeling with a survey of published shape and flow data in ciliates. Model predictions and empirical observations show small variations in feeding rates in favor of either motile or sessile cells. Case-specific variations notwithstanding, our overarching analysis shows that flow physics imposes no constraint on the feeding rates that are achievable by the swimming versus sessile strategies – they can both be equally competitive in transporting nutrients and wastes to and from the cell surface within flow regimes typically experienced by ciliates. Our findings help resolve a long-standing dilemma of which strategy is hydrodynamically optimal and explain patterns occurring in natural communities that alternate between free swimming and temporary attachments. Importantly, our findings indicate that the evolutionary pressures that shaped these strategies acted in concert with, not against, flow physics.

## Introduction

The dense and soluble nature of water allows nutrients necessary for survival to surround small organisms living in both fresh and marine ecosystems [1]. However, the acquisition of these nutrients, either dissolved or particulate, is often challenging because they are frequently dilute or located within sparsely distributed patches [2–5]. Small, single-celled protists near the base of aquatic food chains have faced an evolutionary choice: either swim and use flows generated by swimming to encounter prey, or attach to a substrate and generate feeding currents from which to extract passing particles. Both “swim” or “stay” solutions occur among species in natural communities [1, 6] and a number of species actively alternate between swimming and attachment [7]. Fig. 1A presents a focused survey of these strategies within a pivotal clade of microorganisms, the Ciliophora.

**Figure 1:**
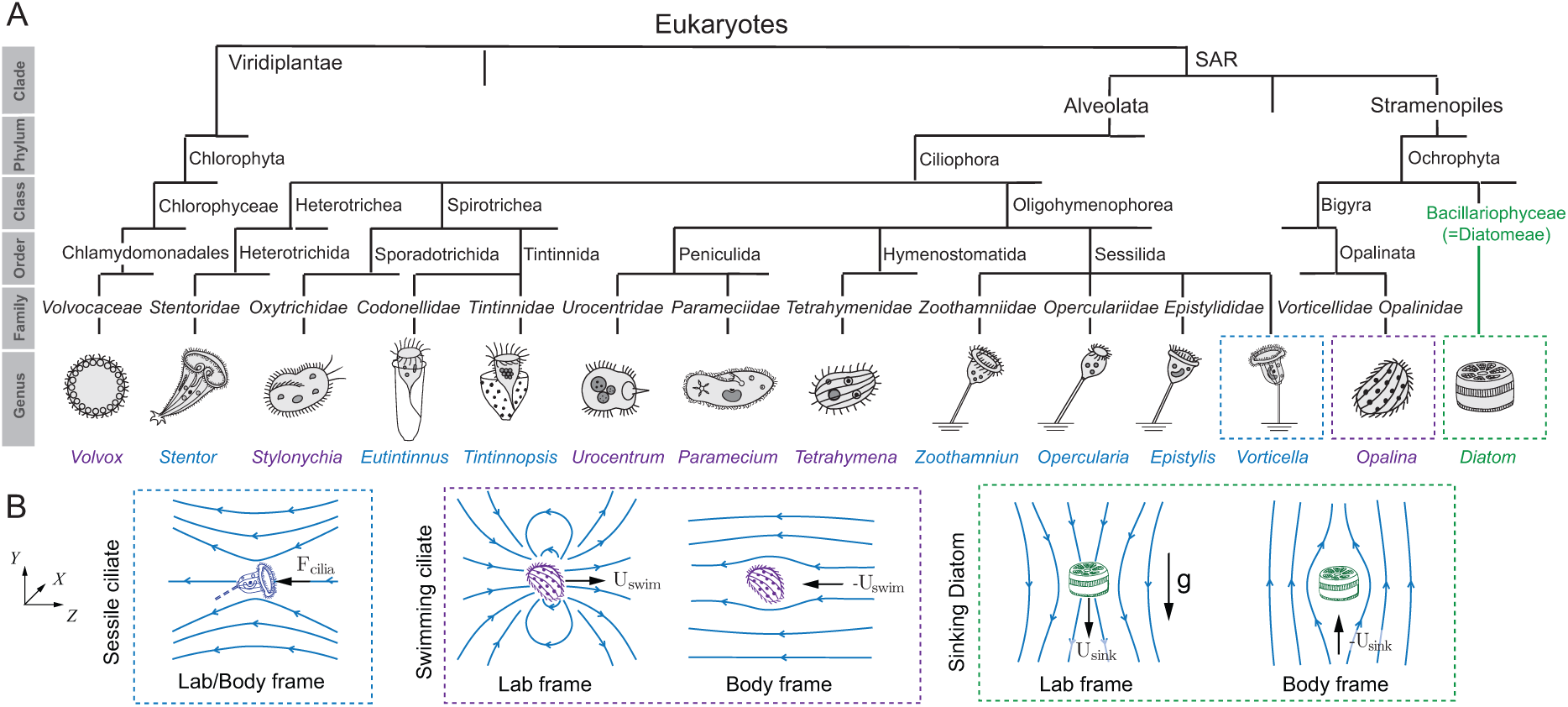
(A) **Phylogenetic tree** showing microorganisms known to feature cilia that generate feeding currents in either sessile (blue) or free swimming (purple) states. The class of diatoms – non-motile cells that sink when experiencing nutrient limitation – is shown for comparison. (B) Flow fields around a sessile ciliate, swimming ciliate, and sinking diatom, in lab and body frame of references. Streamlines are shown in blue in lab frame (*X, Y, Z*).

The “swim” or “stay” strategies shape material transport through this essential link in aquatic trophic systems, thus affecting not only the fitness of these microorganisms, [8–11] but also impacting global biogeochemical cycles and the food web chain [12–15]. Therefore, understanding the flow physics underlying the exchange of nutrients and wastes at this scale is important across disparate fields of the life sciences, from evolutionary biology [16, 17] to ecosystem ecology [18, 19].

It has been generally appreciated that microorganisms, swimming or tethered, manipulate the fluid environment to maintain a sufficient turnover rate of nutrients and metabolites, unattainable by diffusive transport alone [10, 16, 20, 21]. However, to date, and with ample experimental [22, 23] and computational [24–26] studies, flow analysis has yielded contradictory results favoring either of the “swim” [24, 25, 27, 28] or “stay” [22, 23] alternatives as optimal nutritional strategies.

If consideration of flow physics clearly favors one of the “swim” [25] or “stay” [22] alternatives, then the existence of both indicates that the evolutionary pressures that led to the abundance of the other strategy had to act against flow physics and the propensity to optimize material transport to and from the cell surface. It would also imply that both solutions cannot occupy the same ecological niche without one of them being seriously disadvantaged. An alternative possibility is that flow physics supports both solutions equally and that the choice of strategy does not compromise material transport to and from the cell surface. For example, in organisms that alternate between swimming and attachment, this transition is often influenced by external environmental conditions, such as ph balance [29], nutrient concentration [30] and prey availability [31], and predator presence [32].

But how can we distinguish between these two hypotheses? Establishing such a distinction is challenging because any attempt at quantifying flows around a specific microorganism [22, 33] inherently accounts for all evolutionary variables that shaped that microorganism and thus fails to provide a general and unbiased mechanistic understanding of the role of flow physics. Mathematical models allow objective comparison of the feeding rates achievable in the attached versus swimming states, while keeping all other variables the same. Surprisingly, besides [25], there is a paucity of mathematical studies that directly address this question. Importantly, results based on any single model naturally depend on the modeling assumptions; thus, any attempt at drawing general conclusions from considering a single organism or mathematical model should be carefully scrutinized.

In this study, we propose a systematic approach to address existing limitations in evaluating the hydrodynamics of the “swim” or “stay” alternatives. Our approach combines a survey of existing experimental observations within the entire Ciliophora clade (Fig. 1) with mathematical models that span the morphology and flow conditions within which all surveyed ciliates fall (Fig. 2). We additionally include a comparison with diatoms to distinguish the effects of relative body motion independent of cilia-driven feeding currents. We find, based on both empirical observations and mathematical models, that encounter rates of the swim and stay strategies converge under realistic conditions and are essentially equivalent within flow regimes typically experienced by ciliates.

**Figure 2:**
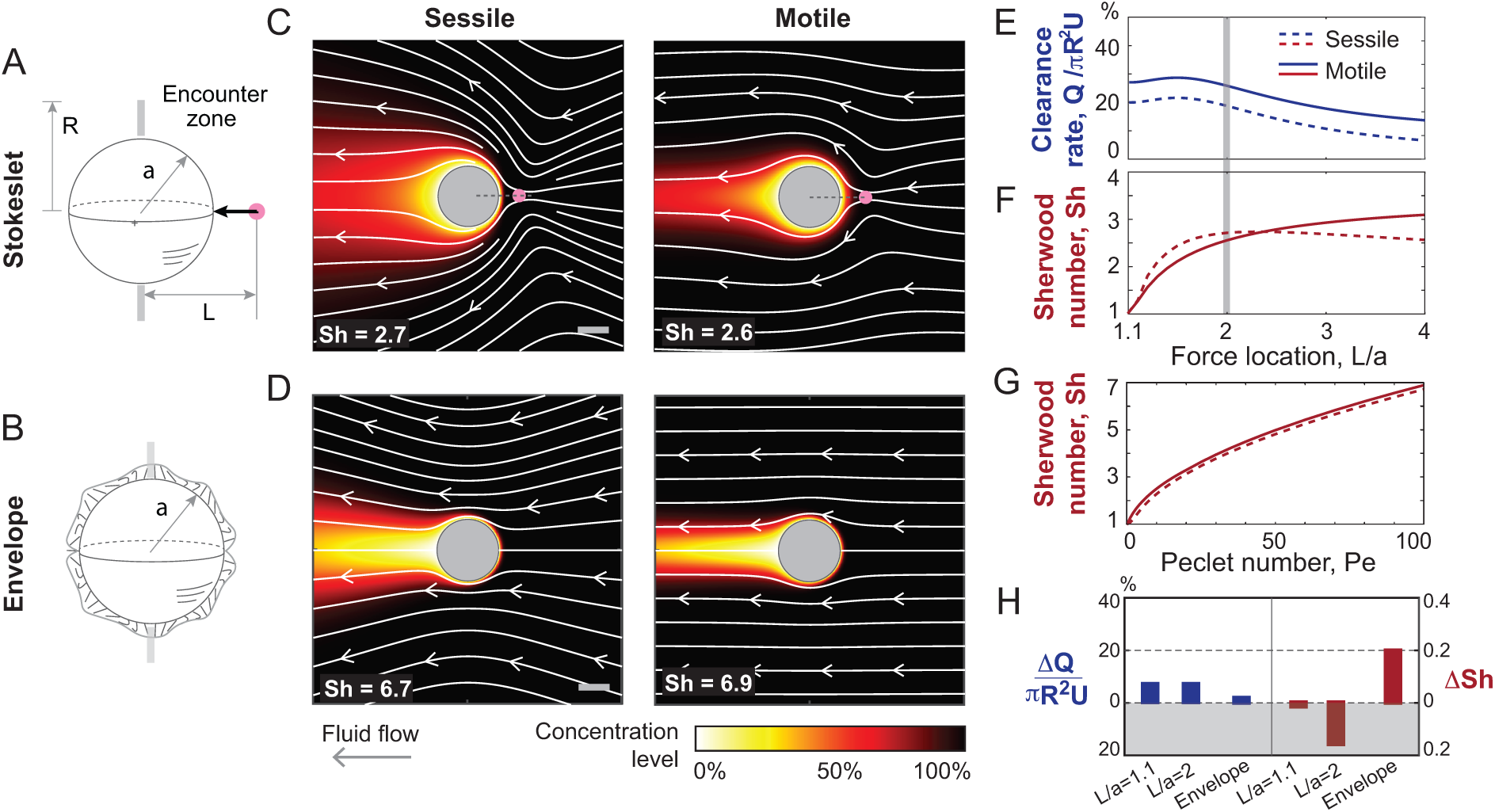
Stokeslet and Envelope models of sessile and motile ciliates. (A) Stokeslet model where ciliary activity is represented by a Stokeslet force *F*_cilia_ is located at a distance (*L a*)*/a* outside the spherical cell surface with no-slip surface velocity. (B)Envelope model where cilia activity is distributed over the entire cell surface with slip surface velocity. (C,D) Fluid streamlines (white) and nutrient concentration fields (colormap) in the sessile and swimming cases. Here, *L/a* = 2, *a* = 1 and *F*_cilia_ is chosen to generate a swimming speed *U* = 2*/*3 in the motile case to ensure consistency with the envelope model. (E) Nutrient uptake in sessile and motile Stokeslet-sphere model based on calculation of clearance rate *Q* of a fluid volume passing through an annular disk of radius *R/a* = 1.1 and Sherwood number Sh. In the latter, Pe is 100. (F) Nutrient uptake in sessile and motile envelope model based on calculation of Sherwood number Sh as a function of Pe. (G) Difference in clearance rate Δ*Q* = *Q*_motile_ *Q*_sessile_ and Sherwood number ΔSh = Δ*I/I*_diffusion_ = Sh_motile_ Sh_sessile_ in the Stokeslet-sphere model for *L/a* = 1.1 and *L/a* = 2 and in the envelope model. In both metrics, the difference is less than 20%: Δ*Q* is less than 20% the advective flux *πR*^2^*U* and Δ*I* is less than 20% of the corresponding diffusive uptake *I*_diffusion_ = 4*πRDC_∞_*. Shaded grey area denotes when the sessile strategy is advantageous.

## Results

Our results are organized around three main themes: (A) comparative analysis of morphologies, size, and fluid flows in sessile and swimming ciliates and sinking diatoms, (B) evaluation of nutrient uptake in mathematical models of sessile and motile feeders spanning the Stokeslet [10, 25, 34] and envelope [35, 36] models and covering the entire range within which all surveyed ciliates fall [37], (C) analysis of biological data in light of model prediction and of asymptotic analysis in the two extremes of diffusion and advection dominant limits.

### Comparative morphometric, phylogenetic and flow data in ciliates and diatoms

We conducted a survey on the morphology, flows, and phylogenetic lineage of ciliates and diatoms, [38–41] (Fig. 1).

Sessile ciliates, such as the *Stentor* [42], *Opercularia* [43, 44], and *Vorticella* [34, 45–48], are characterized by a ciliary crown, where the motion of beating cilia entrains fluid toward the cell. The cell body and ciliary crown are positioned away from the surface they live upon, usually with a stalk, to minimize the effect of that surface on slowing down the cilia-driven microcurrents [21, 34, 49, 50]. Further, to avoid generating recirculating microcurrents and reduce reprocessing of depleted water [43, 47, 51], sessile ciliates actively regulate their orientation to feed at an angle relative to the sub-stratum [34]. At optimal inclination, the effective cilia-generated force is nearly parallel to the bounding substrate and creates quasi-unidirectional flows that drive nutrients and particles past the cell feeding apparatus [34, 52, 53]. In motile ciliates, such as the *Paramecium* and *Volvox*, the surface of the organism is often entirely covered with cilia that beat in a coordinated manner and power the organism to swim through the surrounding fluid [54–57]. Diatoms lack motility apparatus and sink by regulating their buoyancy [20, 58, 59] (Fig. 1B).

Empirical flow measurements around sessile [21, 44, 48, 52, 60, 61] and motile [62, 63] ciliates are sparse. Here, we collected morphometric and flow data from published work covering ten species of sessile ciliates [34, 42, 43, 45, 47, 48], ten species of swimming ciliates [54, 55, 57], and seven species of diatoms [58, 59]. A summary of the ranges of sizes and characteristic speeds are reported in Table 1 and Fig. S1; detailed measurements are listed in a supplemental data file. Size is represented by the volume-equivalent spherical radius *a* (Fig. S2). The characteristic speeds *U* for sessile ciliates are based on the maximal flow speeds measured near the ciliary crown. For swimming ciliates and sinking diatoms, we collected measured swimming and sinking speeds, which, given the no-slip boundary condition in this viscous regime [56, 64], also represent flow speeds near the surface of these microorganisms.

**Table 1:** Survey of size *a* and flow measurements *U* in sessile and swimming ciliates and sinking diatoms (Table S6). Size *a* is calculated using the volume-equivalent spherical radius (Fig. S7). Corresponding ranges of Pe numbers are based on the diffusivity of oxygen, *D* = 10*^−^*^9^ m^2^*·*s*^−^*^1^, live bacteria, *D* = 4 *×* 10*^−^*^10^, m^2^*·*s*^−^*^1^), and dead bacteria *D* = 2*×*10 m *·*s .

| | Empirical measurements | | Péclet number, $\text{Pe}$ | | |
| --- | --- | --- | --- | --- | --- |
| | microorganism size<br>$a$ ($\mu\text{m}$) | characteristic speed<br>$U$ ($\mu\text{m} \cdot \text{s}^{-1}$) | oxygen diffusivity | live bacteria diffusivity | dead bacteria diffusivity |
| Sessile ciliates | 15–60 | 50–2500 | 1–80 | 2–210 | $(5–400) \times 10^3$ |
| Swimming ciliates | 15–180 | 50–3200 | 3–160 | 8–390 | $(17–800) \times 10^3$ |
| Sinking diatoms | 10–120 | 40–210 | 0.4–23 | – | – |

Phylogenetically, all surveyed microorganisms, except the *Volvox*, belong to the SAR supergroup, encompassing the Stramenopiles, Alveolates, and Rhizaria clades (Fig. 1). The Rhizaria clade is not represented in our survey because it mostly consists of ameboids, while its flagellates have complex and functionally-ambiguous morphologies that do not fit in the present analysis [38, 41]. *Volvox*, the only multicellular microorganism listed in Fig. 1, is an algae that belongs to the Viridiplantae clade. Diatoms evolved from the same SAR supergroup as the majority of unicellular ciliates, but without the ciliary motility apparatus, and while early ciliates date back to about 700 million years [65], diatoms appeared later, about 200 million years ago [66–68]. Diatoms generally exist in a suspended state and sink under low nutrient conditions [20, 59, 69]. Of the twelve ciliates listed in Fig. 1, many transition during their lifecycle between sessile and free swimming states [23, 70]. *Stentors* become rounder when swimming [71].

The microcurrents generated by these ciliates improve solute transport to and from the surface of the microorganism. For a characteristic microcurrent of speed *U* = 100 *µ*m *·* s*^−^*^1^, small molecules and particles would be transported over a characteristic distance *a* = 100 *µ*m in approximately *a/U* = 1 s. In contrast, the same substance transported by diffusion alone takes a considerably longer time to traverse the same distance. For example, diffusive transport of oxygen and small molecules, with diffusivities that are in the order of *D* = 10*^−^*^9^ m^2^ *·* s*^−^*^1^, takes about *a*^2^*/D* = 10 s, while live and dead bacterial particles with respective diffusivity *D* = 4 *×* 10*^−^*^10^ m^2^ *·* s*^−^*^1^ and *D* = 2*×*10*^−^*^13^ m^2^ *·*s*^−^*^1^ [72] take about *a*^2^*/D* = 25 s and 10000 s, respectively. The ratio of diffusive *a*^2^*/D* to advective *a/U* timescales defines the Péclet number, Pe = *aU/D*. For Pe *≪* 1, mass transport is controlled by molecular diffusion. For the microorganisms that we surveyed, we obtain Pe ranging from nearly 0 to as large as 10^3^ *−*10^5^ depending on the nutrient diffusivity (Table 1). This dimensional analysis suggests that the flows generated by the microorganisms substantially enhance the transport of solutes to and from their surface, and while it clearly shows that diatoms typically occupy a smaller range of Pe numbers, this analysis does not reveal a clear distinction between sessile and swimming ciliates. To further explore such distinction, if present, and to assess whether ciliates are disadvantaged by flow physics in their attached state compared to their swimming state as suggested in [25], we developed mathematical models that allow for an unbiased comparison between these two states.

### Mathematical modeling of fluid flows and nutrient uptake

To quantify and compare nutrient uptake across microorganisms, we approximated the cell body by a sphere of radius *a*, as typically done in modeling sessile and swimming ciliates [25, 35, 36, 73] and sinking diatoms [10, 20, 74] (Figs. 2).

The fluid velocity **u** around the sphere is governed by the incompressible Stokes equations, *−∇p* + *η∇*^2^**u** = 0 and *∇ ·* **u** = 0, where *p* is the pressure field and *η* is viscosity. We solved these equations in spherical coordinates (*r, θ, ϕ*), considering axisymmetry in *ϕ* and proper boundary conditions. In the motile case, we solved for the fluid velocity field **u** in body frame by superimposing a uniform flow of speed *U* equal to the swimming speed past the sphere; we calculated the value of *U* from force balance considerations [25, 75] (see SI for details).

We solved the Stokes equations for two models of cilia activity: cilia represented as a Stokeslet force *F*_cilia_ placed at a distance *L* and pointing towards the center of the sphere and no-slip velocity at the spherical surface [25, 76–78] (Fig. 2A), and densely-packed cilia defining an envelope model with a slip velocity **u***|_r_*_=_*_a_* = *U* sin *θ* at the spherical surface where all cilia exert tangential forces pointing from one end of the sphere to the opposite end [24, 35, 36] (Fig. 2B). Detailed expressions of the flow fields and governing equations in both models are included in the SI. In dimensionless form, we set the cell’s length scale *a* = 1 and tangential velocity scale *U* = 1 in the envelope model, and we set the ciliary force *F*_cilia_ in the Stokeslet model to produce the same swimming speed (*U* = 2*/*3) as in the envelope model when the sphere is motile.

To evaluate the steady-state concentration of dissolved nutrients around the cell surface, we numerically solved the dimensionless advection-diffusion equation Pe **u** *·∇C* = Δ*C* in the context of the Stokeslet and envelope models. Here, the advective and diffusive rates of change of the nutrient concentration field *C*, normalized by its far-field value *C_∞_*, are given by Pe **u** *· ∇C* and Δ*C*, respectively, with *∇C* the concentration gradient. At the surface of the sphere, the concentration is set to zero to reflect that nutrient absorption at the surface of the microorganism greatly exceeds transport rates of molecular diffusion [79–81].

In Fig. 2C,D, flow streamlines (white) and concentration fields (colormap at Pe = 100) are shown in the Stokeslet and envelope models. In the sessile sphere, ciliary flows drive fresh nutrient concentration from the far-field towards the ciliated surface. These fresh nutrients thin the concentration boundary layer at the leading surface of the sphere, where typically the cytostome or feeding apparatus is found in sessile ciliates, with a trailing plume or “tail” of nutrient depletion. Similar concentration fields are obtained in the swimming case, albeit with narrower trailing plume.

To assess the effects of these cilia-generated flows on the transport of nutrients to the cell surface, we used two common metrics of feeding. First, we quantified fluid flux or clearance rate *Q* through an encounter zone near the organism’s oral surface [17, 22, 34]. Namely, following [25], we defined the clearance rate *Q* = *−*2*π* ∫*^R^*_*a*_ **u** *·* **e** *| rdr*, normalized by the advective flux *πR*^2^*U*, over an annular encounter zone of radius *R* extending radially away from the cell surface (Fig. 2A). Second, we quantified the concentration flux of dissolved nutrients at the cell surface [10, 24, 36]. To this end, we integrated the inward concentration flux *I* = ∫*_S_ D∇C ·* **n**^*dS*, normalized by the diffusive nutrient uptake *I*_diffusion_ = 4*πRDC_∞_* to get the Sherwood number Sh = *I/I*_diffusion_. We applied both metric to each of the Stokeslet and envelope models.

In Fig. 2E, we report the clearance rate *Q* in the context of the Stokeslet model as a function of the ciliary force location *L/a* for a small annular encounter zone of radius *R* = 1.1*a* extending away from the cell surface. Swimming is always more beneficial. However, the increase in clearance rate due to swimming is less than 10%. This is in contrast to the several fold advantage obtained in [25] for *L* = 4*a* and *R* = 10*a*. (results of [25] are reproduced in Fig. S3). We employed the same metric *Q* in the envelope model and found that motility is also more advantageous, albeit at less than 5% benefit (Fig. 2H).

A few comments on the choice of the size of the encounter zone are in order. Nutrient encounter and feeding in ciliates occur near the leading edge of the ciliary band [82–85]. Cilia are typically of the order of 10 microns in length, and the cell body of a ciliate is typically in the range of 10-1000 microns. We chose *R* = 1.1*a* indicating encounter within an annular protrusion extending 10% beyond the body radius because it falls within the biological range and because a larger encounter zone would induce additional drag on the body that needs to be accounted for in the model. In contrast, [25] chose an encounter zone extending up to 900% the body radius, without accounting for the drag that such a large collection area would add to a swimming body. This also exceeds biological considerations in most ciliates and flagellates, even in *Choanoflagellates* [86] and *Chlamydomonas* [86], where the flagellum length could be up to six times the cell radius.

In Fig. 2F and G, we report the Sh number based on the Stokelet and envelope models, respectively. In the Stokeslet model (Fig. 2F), sessile spheres do better when the cilia force is close to the cell surface (*L − a*)*/a* ⪅ 1.25. In the envelope model (Fig. 2G), motile spheres do slightly better for all Pe⪅ 100. The difference ΔSh between the sessile and motile spheres favors, by less than 20%, the sessile strategy in the Stokeslet model and the swimming strategy in the envelope model (Fig. 2H).

Comparing Sh between the Stokeslet and envelope models (Fig. 2C and D), we found that, at Pe = 100, Sh = 2.7 (sessile) and 2.6 (motile) in the Stokeslet model compared to Sh = 6.7 (sessile) and 6.9 (motile) in the envelope model. This is over a two-fold enhancement in nutrient uptake at the same swimming speed *U* = 2*/*3 simply by distributing the ciliary force over the entire surface of the cell! Indeed, this improvement occurs because the ciliary motion in the envelope model significantly thins the concentration boundary layer along the entire cell surface as opposed to only near where the cilia force is concentrated in the Stokeslet model.

In our survey of sessile and motile ciliates (Fig. 1), cilia are clearly distributed over the cell surface. Thus, we next explored in the context of the envelope model the behavior of the Sh number across a range of Pe values that reflect empirical values experienced by the surveyed ciliates (Table 1).

### Linking model prediction to biological data

We numerically computed the Sherwood number for a range of Pe *∈* [0, 1000] for the sessile and motile spheres, and, to complete this analysis, we calculated the Sh number around a sinking sphere. Numerical predictions (Fig. 3A, solid lines, log-log scale) show that at small Pe, swimming is more advantageous than attachment; in fact, any motion, even sinking, is better than no motion at all [16]. However, at larger Pe, there is no distinction in Sh number between the sessile and motile sphere.

**Figure 3:**
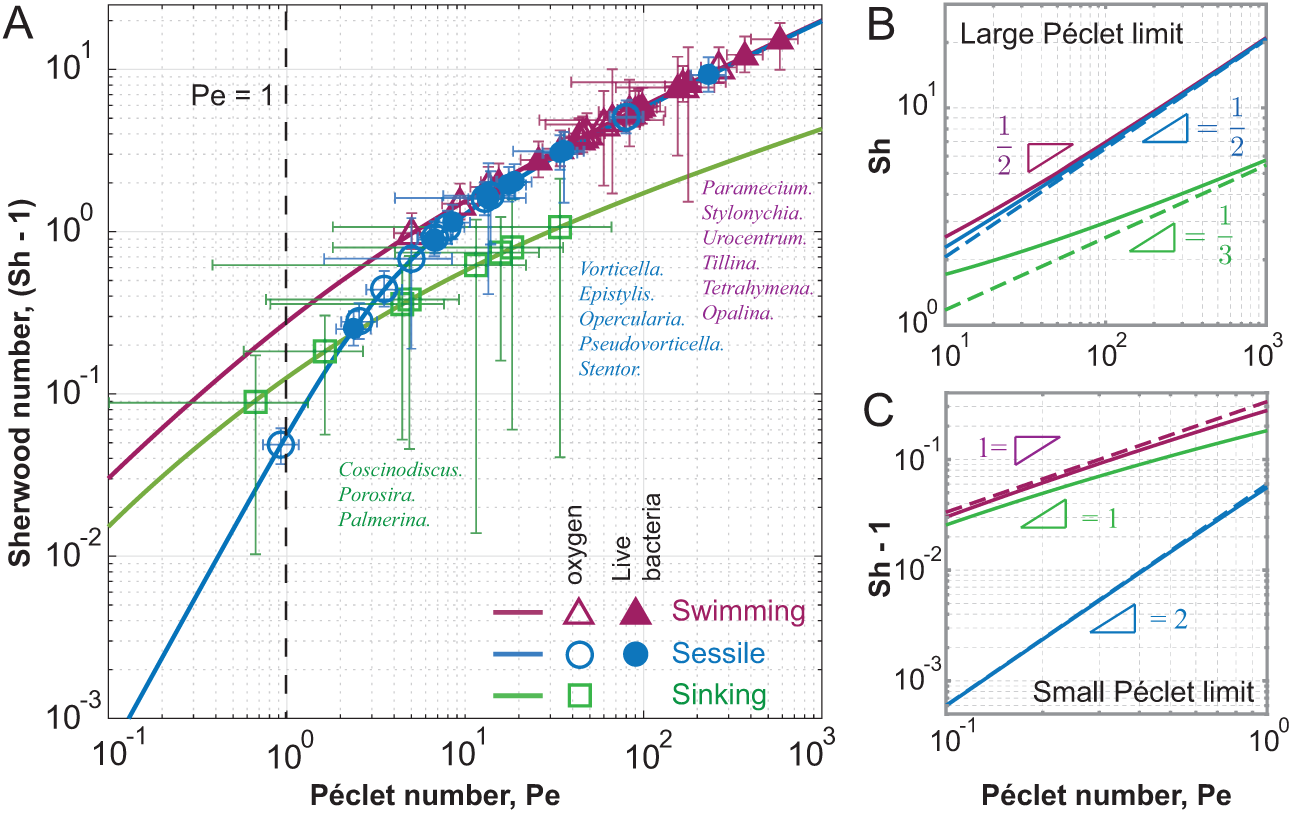
**Sherwood number versus Péclet number** for the sinking (green) diatom and the swimming (purple) and sessile (blue) ciliates based on the envelope model. (A) Shifted Sherwood number (Sh - 1) versus Péclet number in the logarithmic scale for a range of Pe from 0 to 1000. Pe numbers associated with experimental observations of diatoms (square), swimming ciliates (triangle), and sessile ciliates (circle) are superimposed. Corresponding Sh numbers are calculated based on the mathematical model. Empty symbols are for oxygen diffusivity *D* = 1 × 10*^−^*^9^m^2^ s*^−^*^1^ and the solid symbols correspond to the diffusivity *D* = 4 × 10*^−^*^10^m^2^ s*^−^*^1^ of live bacteria [72]. (B-C) Asymptotic analysis (dashed lines) of Sherwood number in the large Péclet limit (B) and small Péclet limit (C).

We next used as input to the sessile, swimming, and sinking sphere models, the Pe numbers obtained from experimental measurements of sessile (blue *⃝*) and swimming (purple *△*) ciliates and sinking diatoms (green □), respectively, and we computed the corresponding values of Sh number (Fig. 3A). Sinking diatoms are characterized by smaller values of Sh number, whereas with increasing Pe, the Sh values of sessile ciliates span similar ranges as those of swimming ciliates.

To complete this analysis, we probed the feeding rates under extreme Péclet limits. We extended the asymptotic scaling analysis developed in [87, 88] and translated to nutrient uptake in sinking diatoms [20] and swimming ciliates [36, 73], to arrive at asymptotic expressions for sessile ciliates in the two limits of small and large Pe,

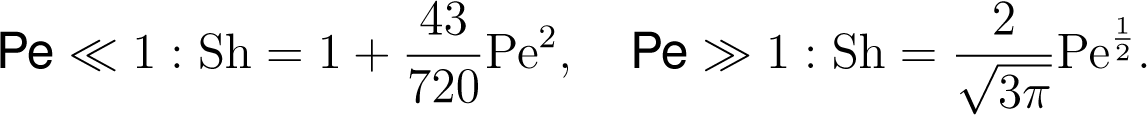

In Fig. 3B and C, we superimposed our asymptotic results, together with the asymptotic results of [20, 36, 73, 87, 88], onto our numerical findings. At small Pe *≪* 1, the Sh numbers for swimming and sinking spheres scale similarly with Pe (Sh *∼* Pe), whereas Sh scales worse (Sh *∼* Pe^2^) for the sessile sphere. Our thorough literature survey indicates, save one, no data points for sessile microorganisms in this limit. At large Pe *≫* 1, the Sh numbers of the sessile and swimming spheres scale similiary with Pe (Sh *∼* Pe 2), whereas the sinking sphere scales worse (Sh *∼* Pe 3). Similar scaling is found in swimming ciliate models [36, 73, 81]. These results confirm that, hydrodynamically, sessile and swimming ciliates are equivalent in the limit of large Pe. When cilia generate strong feeding currents that drive nutrients and particulates toward the cell body, attached microorganisms can be equally competitive with motile microorganisms that swim to feed.

## Discussion

We contributed a comprehensive methodology for evaluating the role of flow physics and comparing feeding rates in motile and sessile ciliates. Our approach combined a survey of previously published empirical measurements of ciliates’ shape and velocity with two mechanistic models of cilia-driven flows (concentrated point force and distributed force density) and two metrics of nutrient uptakes (clearance rate and Sherwood number) in attached and swimming ciliates. The concentrated versus distributed ciliary force models form two extreme limits within which all surveyed ciliates fall. Clearance rate measures advective material transport through an encounter zone, which is independent of Pe; Sh number accounts for both diffusive and advective transport and varies with Pe.

The difference in feeding rates between the sessile and motile strategies depended on the choice of model, model parameters, and feeding metric (Fig. 2).

In the context of the concentrated force model and considering clearance rate as a metric for feeding, we found that it is better to swim than to attach, but these advantages are modest (less than 20%) under justifiable conditions of ciliary force placement and encounter zone close to the cell surface. In [25], several fold improvement were reported for swimming using the same model and feeding metric but questionable parameter values - clearance rates were computed through an encounter zone that extended up to ten body lengths away from the cell surface without accounting for the effect that such an extensive collection surface would have on drag generation during swimming [25]. We showed that for a small encounter zone that justifies omission of these drag forces, the improvement in clearance rate during swimming is much smaller than predicted in [25].

Surprisingly, using the same concentrated force model, we found that attachment improves nutrient uptake when considering concentration of dissolved nutrients at Pe = 100 and measuring the Sherwood number associated with nutrient uptake over the entire cell surface. Again, the improvement is modest (Fig. 2H). Taken together, these results show that in the same model, two different feeding metrics favor different strategies, albeit at a slim advantage of less than 20% in favor of either swimming or attachment.

When distributing the ciliary force over the entire cell surface, we found, using either metric, that swimming is more beneficial by a very small margin for Pe *≤* 100 (Fig. 2G). Interestingly, the difference in Sh number between swimming and attached cells decreases at larger Pe values (Fig. 3A), and in the asymptotic limit of Pe*≫* 1, Sh scales similarly with Pe (Sh *∝* Pe^1^*^/^*^2^) for both swimming and sessile cells (Fig. 3A and B). That is, at large Pe, material transport to and from the cell surface is not compromised by the choice of strategy.

From our survey of previously-published empirical measurements of ciliates’ shape and velocity (Fig. 1), we extracted biologically-relevant ranges of Pe values (Table 1) and combined these empirical observations with model predictions (Fig. 3A). We found significant overlap in Sh number between sessile and motile organisms at a wide range of representative Pe values. These findings clearly show that both attachment and free swimming can lead to similar nutrient acquisition within a wide range of flows and Péclet values typically experienced by ciliates.

This study provides a fresh perspective on evaluating the role of flow physics in the feeding strategies of microorganisms. Prior methods in support of either the motile or sessile strategies as optimal drew general conclusions from focused analyses. Support for swimming came principally from flow-based models of idealized organisms propelling themselves through water [24, 25, 27, 28]. Support for maximum feeding by attached protists came from empirical measurements of prey removal by swimming versus attached individuals [22, 23]. Our approach shows that, while feeding rates may vary between organisms and mathematical models, given a cellular (ciliary) machinery that allows microorganisms to manipulate the surrounding fluid and generate flows, flow physics itself imposes no constraint on what is achievable by the swimming versus sessile strategies – they can both be equally competitive in transporting nutrients and wastes to and from the cell surface in the large Pe limit where nutrient advection is dominant. Our findings suggest that the choice of feeding strategy was likely influenced by evolutionary, ecological or behavioral variables other than flow physics, such as metabolic or sensory requirements [89–91], predator avoidance [92], symbiotic relations [10], and nutrient availability or environmental turbulence [26, 92, 93].

Along with assessing feeding rates in motile versus sessile strategies, our analysis revealed interesting “design” principles for maximizing nutrient uptake by distributing ciliary activity over the entire cell surface (Fig. 2). This design thins the nutrient-depletion boundary layer at the surface of the cell where absorption occurs: for the same overall swimming speed, distributing ciliary activity over the cell surface improves nutrient uptake by over two fold compared to when the ciliary force is concentrated at one location (Fig. 2). Indeed, cilia are often distributed over a portion or entire cell surface in sessile and motile ciliates, with some variability in cilia distribution and cell surface fraction where prey is intercepted (Fig. 1). To account for such variability, we computed the flow and concentration fields under various perturbations to cilia coverage and surface fraction where absorption takes place (Fig. 4). For each perturbation, we calculated the Sh number in the form of a percentage of that corresponding to full cilia coverage and absorption over the entire surface. We found small differences in Sh numbers between the sessile and motile spheres. Our findings – that the motile and sessile strategies are equivalent in terms of material transport to the cell surface – are thus robust to cilia perturbations. Additionally, we found that, for a given cilia coverage, nutrient uptake is maximized when the absorption surface coincides with the cilia coverage area. This design – cilia collocated with the cell feeding apparatus – is abundant in sessile protists (Fig. 1). Our findings open new venues for investigating the functional advantages of optimal cilia designs (cilia number and distribution) that maximize not only locomotion performance [37] but also feeding rates and for evaluating the interplay between cell design and feeding strategies (sessile versus motile) both at the unicellular [94] and multicellular levels [95]. These future directions will enrich our understanding of the complexity of feeding strategies in ciliates and how strategy and design have evolved to provide behavioral advantages to these microbes.

**Figure 4:**
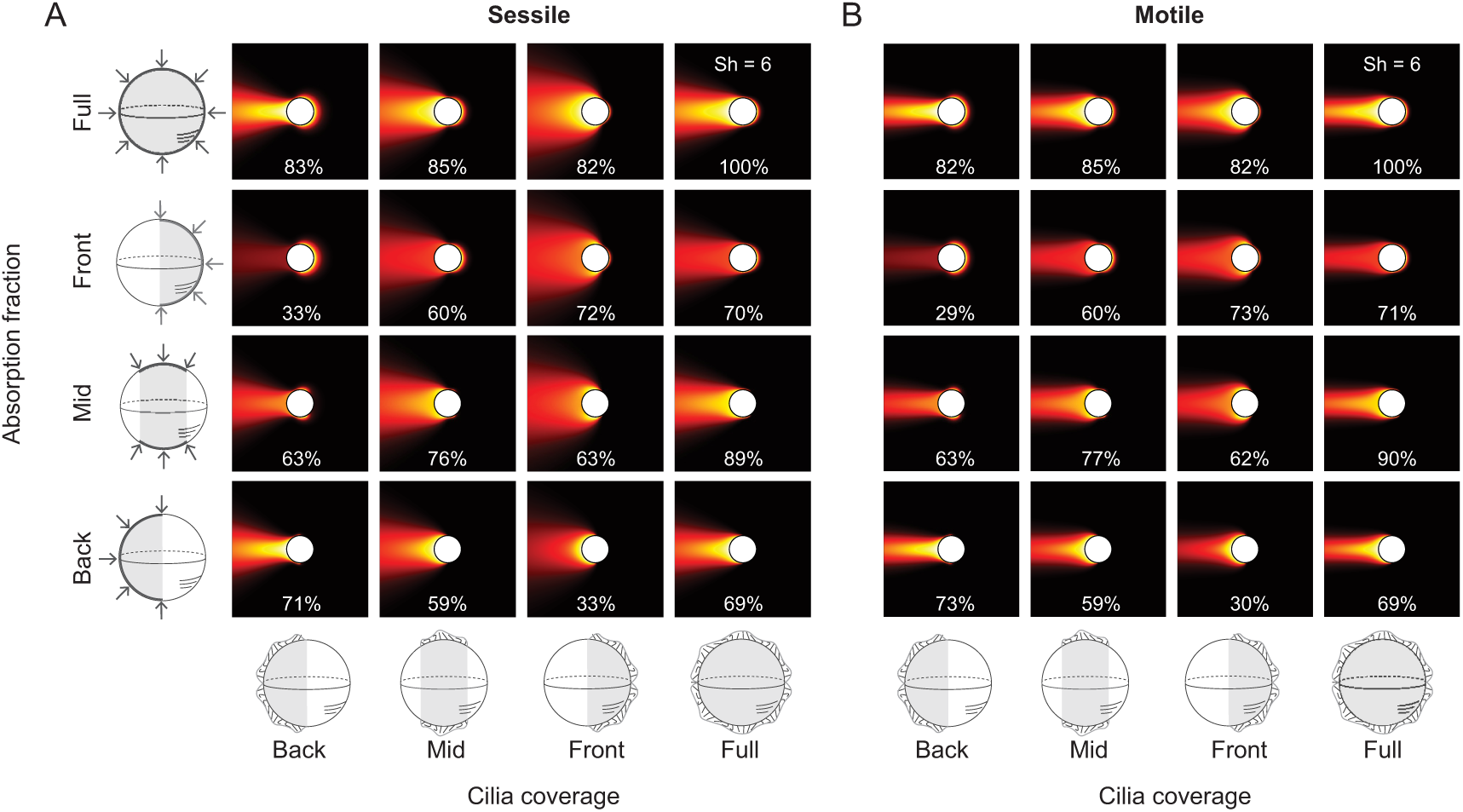
Robustness to variations in cilia coverage and absorption fraction. We considered a 50% cilia coverage and 50% absorption fraction located at back, middle, and front of the (A) sessile and (B) motile sphere. Concentration fields and Sherwood numbers with 100% cilia coverage and absorption area are shown in the top right corner. In all other cases, the Sh number is reported as a percentage of the full coverage/absorption case.

## Acknowledgment

EK acknowledges support from the University of Southern California (for PhD student JL) and support from the Office of Naval Research (ONR) Grants N00014-22-1-2655, N00014-19-1-2035, N00014-17-1-2062, and N00014-14-1-0421; the National Science Foundation (NSF) Grants RAISE IOS-2034043, CBET-2100209, and INSPIRE MCB-1608744; the National Institutes of Health (NIH) Grant R01 HL 153622-01A1; and the Army Research Office (ARO) Grant W911NF-16-1-0074.

## Notes

### Competing Interest Statement

The authors have declared no competing interest.

### Summary of Updates

Revised main text and SI following acceptance to eLife

