## Supplemental Material and Figures for "Feeding Rates in Sessile versus Motile Ciliates are Hydrodynamically Equivalent"

### 1 Collection of Published Biological Data

#### Kinematic measurements

We collected from published literature measurement data of microorganisms, including size, swimming speed of motile organisms, and surrounding flow speed of sessile organisms. For modeling and further analyzing, we simplified the shape of those organisms as a sphere with equivalent sphere diameters. Based on each organism's coarse morphology (Fig. 1.A), we first approximated these morphologies into regular shapes (Fig. 1.B), then, by calculating the volume of those shapes, we solved for the radius $a$ of a model sphere that has the same volume. The data is included in the attached excel sheet.

Using the raw data, we plot $U/a$ in Fig. 2.A in regular scale and in Fig. 2.B in logarithmic scale using the equivalent spherical radius $a$ as the length scale, the swimming speed as the

*

velocity scale $U$ for the motile organism, and the maximum observed flow speed as the velocity scale $U$ for sessile organisms. In addition, we plot $Ua$ in Fig. 2.C, representing advection strength.

#### Phylogenetic information

For each species for which we collected morphometric and flow kinematic data, we traced it back along the evolutionary tree. The diatoms, swimming, and sessile ciliates were traced back to their common Eukaryotic ancestor. From that common ancestor, various clades evolved [1], [2], [3], [4], including the supergroup SAR from which ciliates descended, and the Viridiplantae group to which green algae like the volvox belong. We combined the taxonomy information of the species collected above to obtain the tree of life diagram in the main text.

### 2 Mathematical Modeling

#### Stokes equations

At the microscopic scale, the Reynolds number Re which quantifies the ratio of inertial to viscous forces is nearly zero. The viscous forces are dominant, and the fluid velocity $\mathbf{u}$ is governed by the incompressible Stokes equation,

$$-\nabla p + \eta \nabla^2 \mathbf{u} = 0, \qquad \nabla \cdot \mathbf{u} = 0, \tag{1}$$

where $p$ is the pressure field and $\eta$ is the dynamic viscosity.

#### Stokeslet model

Following [5, 6], we represented the ciliary activity as a point force $\mathbf{F} = -F_{\text{cilia}}\mathbf{e}_z$ located at $L\mathbf{e}_z$, at a distance $L$ from the center of the sphere (Fig. 3). For the motile sphere, the flow field is obtained by superposition of two solutions of the Stokes equation, to obtain a solution with no slip boundary condition at the sphere's surface: $\mathbf{u} = \mathbf{u}_1 + \mathbf{u}_2$, where $\mathbf{u}_1 = \mathcal{G}_o \cdot \mathbf{F}$ and $\mathcal{G}_o$ is Green's tensor due to a point force near a rigid sphere [7, 8] and $\mathbf{u}_2$ is the solution due to uniform flow $-U\mathbf{e}_z$ pass a rigid sphere, expressed in the sphere's frame of reference [5, 8].

In the Stokes regime, force balance on the swimming sphere is given by

$$\mathbf{0} = \mathbf{T} + \mathbf{K} + \mathbf{D}, \tag{2}$$

where $\mathbf{T} = -\mathbf{F}$ is the thrust force generated by the flagella (Stokeslet) in the swimming direction $\mathbf{e}_z$, $\mathbf{D}$ is the drag force due to a moving sphere in fluid with speed $U$, and $\mathbf{K}$ is the hydrodynamic force acting on the sphere by the flow generated by the point force $\mathbf{F}$. Evaluating these terms leads to a scalar equation in the $\mathbf{e}_z$-direction that can be solved to obtain the swimming speed

$$U = \frac{F_{\text{cilia}}}{6\pi\mu a}\left(1 - \frac{3a}{2L} + \frac{a^3}{2L^3}\right). \tag{3}$$

For a sessile sphere, The force balance on the sphere is given by:

$$\mathbf{0} = \mathbf{F}_{\text{tether}} + \mathbf{T} + \mathbf{K}, \tag{4}$$

where $\mathbf{T} = -\mathbf{F}$ and $\mathbf{K}$ are defined as above. This equation can be solved to obtain the force $\mathbf{F}_{\text{tether}}$ provided by a tether to prevent the sphere from moving in the $\mathbf{e}_z$ direction. Namely,

$$\mathbf{F}_{\text{tether}} = \left(1 - \frac{3a}{2L} + \frac{a^3}{2L^3}\right) F_{\text{cilia}}\mathbf{e}_z, \tag{5}$$

To non-dimensional these solutions in (3) and (5), we chose $a$ as the characteristic length scale and $\mathcal{U} = F_{\text{cilia}}/(8\pi\mu a)$ as the characteristic velocity scale. Results based on our implementation are shown in Fig. 3 and are consistent with the results of [5].

### Envelope model

We reproduced the general solution of (1) subject to arbitrary slip velocity at the surface of the sphere and proper decay at infinity. The solution is given in terms of the radial distance $r$ from the center of the sphere and an angular variable $\mu = \cos\theta$, in the form of an expansion in Legendre polynomials $P_n(\mu)$ [9–11].

We considered the "treadmill" slip velocity at the surface of the sphere

$$u_\theta|_{r=a} = B\sqrt{1-\mu^2}, \qquad u_r|_{r=a} = 0, \tag{6}$$

where $B$ is a constant parameter that defines a velocity scale $\mathcal{U}$. In this case, the solution simplifies significantly. In Table 1, we list mathematical expressions for the boundary conditions, fluid velocity field, pressure field, forces acting on the sphere, hydrodynamic power, and swimming speed for freely moving ciliated sphere. Results based on our analysis are consistent with the results of [10] for a swimming sphere.

### Clearance rate

The clearance rate is calculated by integrating the $z$-component of the flow velocity $u_z$ past an annular disk around the equator of the sphere (Fig. 3). Normalizing the result by a flux $\pi R^2 U$ of a uniform flow $U$ through a disk of area $\pi R^2$, we get the clearance rate (with minus sign to indicate positive clearance value)

$$Q = -\frac{2}{R^2 U}\int_a^R u_z|_{z=0} r dr. \tag{7}$$

When applied to the Stokeslet model, this clearance rate is consistent with that in [5] (Fig. 3). By applying it to the fluid solution in the envelope model (Table 1), we can compare feeding rates across the two models ciliary models which reflect different distributions of the ciliary force – concentrated in the Stokeslet model versus fully distributed in the envelope model.

### Advection-diffusion equation

To determine the effect of the advective currents generated by the attached and freely-swimming ciliated sphere on the concentration distribution around the sphere, we considered the advection-diffusion equation for the steady-state concentration $C$ of nutrients around the spherical surface

$$\mathbf{u} \cdot \nabla C = D \Delta C. \tag{8}$$

Here, $\mathbf{u} \cdot \nabla C$ and $D\Delta C$ are, respectively, the advective and diffusive rates of change of the nutrient concentration field $C$. Considering the absorption of nutrients is limited by number of surface receptors, we assume nutrient concentration on sphere surface is constant [12, 13]. Thus, by removing the constant from the background concentration, the concentration at the spherical surface can be considered zero [14, 15]. The boundary conditions, for both the attached and free-swimming spheres, are given by

$$C(\mu)|_{r=a=1} = 0, \qquad C(\mu)|_{r\to\infty} = C_\infty. \tag{9}$$

For non-dimensionalization, we scale the concentration using $c = (C_\infty - C)/C_\infty$ assuming zero concentration at an infinite distance and uniform concentration on the sphere's surface. By choosing characteristic velocity scale $\mathcal{U}$ and length scale $a$, we obtain the dimensionless governing equation and boundary conditions

$$\mathrm{Pe}\mathbf{u} \cdot \nabla c = \nabla^2, \quad c(r = 1) = 1, \quad c(r \to \infty) = 0, \tag{10}$$

where $\mathrm{Pe} = a\mathcal{U}/D$.

**Sherwood number.** By Fick's law, a gradient in concentration yields a flux. The nutrient uptake rate is the area integral of the flux over the spherical surface $I = -\oint \hat{\mathbf{n}} \cdot (-D\nabla C)\mathrm{d}S$, where $\mathrm{d}S = 2\pi R^2 \sin\theta \mathrm{d}\theta$ is the element of surface area of the sphere. This sign convention is such that the concentration flux is positive if the sphere takes up nutrients. In the absence of microcurrents, the concentration is governed by diffusion only. The steady-state concentration obtained by solving the diffusion equation $\nabla^2 c = 0$ internally bounded by a sphere is given by $c(r) = 1/r$ in non-dimensional form; for which, $C(r) = C_\infty(1 - a/r)$. The steady-state inward current due to molecular diffusion is given by $I_{\text{diffusion}} = 4\pi a D C_\infty$.

For diffusion coupled with advective microcurrents, the nutrient uptake is quantified by the Sherwood number Sh, which is equivalent to a dimensionless nutrient uptake, where $I$ is scaled by $I_{\text{diffusion}}$. The dimensionless form of Sherwood number is

$$\mathrm{Sh} = -\frac{1}{2}\int_{-1}^{1} \nabla c \cdot \mathbf{e}_r\big|_{r=1}\, d\mu. \tag{11}$$

Eq. (11) provides an alternative metric for evaluating nutrient uptake to the one in (7). By applying (7) and (11) to the Stokeslet and envelope models, we are able to compare nutrient uptake across two ciliary models that reflect different distributions of ciliary forces – concentrated versus fully distributed force – using two different metrics. Having two metrics – flow rate (7) and Sherwood number (11) – allows us to test robustness of the results to the choice of metric. For example, when applied to the Stokeslet model, (11) allows us to test the robustness of the results in [5] to the choice of metric, concluding that their results are not universal.

**Numerical methods.** Given the symmetry of all fluid fields under consideration, we solve for the corresponding concentration fields in spherical coordinates $(r, \theta, \phi)$ with $\phi$ axis-symmetry. To solve for the Stokeslet model, we use finite difference method on a two-dimensional mesh $(r, \theta)$ that is stretched to ensure a denser grid near the sphere and a sparser grid further away [16]. And for solving the envelope model, we use the Legendre spectral method [15], which uses Legendre polynomials as the spectrum basis in $\theta$ dimension and finite difference in $r$ dimension with a stretched mesh.

### Asymptotic analysis

For analyzing the advection-diffusion equation under extreme Péclet number $\mathrm{Pe} \ll 1$ and $\mathrm{Pe} \gg 1$, we use the approach employed in [17, 18] for a rigid sphere and in [14] for the spherical envelope model.

**Small Pe.** At small Péclet number, we expand the concentration field as

$$c = \mathrm{Pe}^0 c_0 + \mathrm{Pe}^1 c_1 + \mathrm{Pe}^2 c_2 + \ldots \tag{12}$$

We substitute the expanded concentration into the dimensionless advection-diffusion equation (10), we arrive at system of equations associated with each order in Pe, $\mathrm{Pe}^0$, $\mathrm{Pe}^1$, $\mathrm{Pe}^2$. At the leading order $\mathrm{Pe}^0$, the solution is simply $c_0 = 1/r$. To find the solution at higher orders, we substitute the velocity field corresponding to each model (see Table.1) and solve for the higher order equations. We solve for the equations associated with the first three order of Pe, the asymptotic solution to $\mathrm{Pe} \ll 1$ for all considered models are in Table.3.

**Large Pe.** For $\mathrm{Pe} \gg 1$, we take the Taylor series expansion of flow field at $r = 1$ and keep only the leading terms

$$\begin{aligned} u_r(r,\mu) &= u_r|_{r=1} + \frac{\partial u_r}{\partial r}\Big|_{r=1}(r-1) + \ldots, & u_\theta(r,\mu) &= u_\theta|_{r=1} + \frac{\partial u_\theta}{\partial r}\Big|_{r=1}(r-1) + \ldots, \\ &= -2(r-1)\mu + \ldots, & &= \sqrt{1-\mu^2} + \ldots. \end{aligned} \tag{13}$$

We define the temporary variable $y = r - 1$ (not to confused with the $y$-coordinated in the inertial $(x, y, z)$ space). The region $y \ll 1$ represents a thin boundary layer around the spherical surface. Since the concentration boundary layer is expected to be thinner as Pe increases, we rescale $r - 1 = y = \mathrm{Pe}^{-m} Y$, where $Y$ is a new variable. By substituting the new variable and the linearized flow field (13) into the advection-diffusion equation (8), and matching order of

$\mathrm{Pe}$ on both sides of the advection-diffusion equation, we obtain $m = 1/2$ for envelope model and $m = 1/3$ for rigid sphere model. By substituting $m$ with keeping only the leading order, we can transfer the partial differential equation into ordinary equation by inducing a new similarity variable [14]. By solving the ODEs, we obtain the asymptotic solutions to $\mathrm{Pe} \gg 1$ for all models listed in Table.3.

Table 1: Envelope model subject to treadmill slip velocity. Mathematical expressions of boundary conditions, fluid velocity field, pressure field, forces acting on the sphere, hydrodynamic power, and speed for each representative case: sessile ciliated sphere, freely swimming ciliated sphere, and sinking (non-ciliated) sphere. All quantities are given in dimensional form in terms of the radial distance $r$ and angular variable $\mu = \cos\theta$.

| | **B.C. at the surface of the sphere** | **B.C. at infinity** |
|---|---|---|
| Sessile ciliated sphere | $u_\theta\|_{r=a} = B\sqrt{1-\mu^2}, \quad u_r\|_{r=a} = 0$ | $\mathbf{u}\|_{r\to\infty} = \mathbf{0}$ |
| Swimming ciliated sphere | $u_\theta\|_{r=a} = B\sqrt{1-\mu^2}, \quad u_r\|_{r=a} = 0$ | $\mathbf{u}\|_{r\to\infty} = -U\mathbf{e}_z$ |
| Sinking (non-ciliated) sphere | $\mathbf{u}\|_{r=a} = 0$ | $\mathbf{u}\|_{r\to\infty} = -U\mathbf{e}_y$ |

| | **Fluid velocity field** |
|---|---|
| Sessile | $u_r(r,\mu) = \left(\frac{a^3}{r^3} - \frac{a}{r}\right) B\mu, \qquad u_\theta(r,\mu) = \frac{1}{2}\left(\frac{a^3}{r^3} + \frac{a}{r}\right) B\sqrt{1-\mu^2}$ |
| Swimming | $u_r(r,\mu) = \left(-\frac{2}{3} + \frac{2a^3}{3r^3}\right) B\mu, \quad u_\theta(r,\mu) = \left(\frac{2}{3} + \frac{a^3}{3r^3}\right) B\sqrt{1-\mu^2},$ |
| Sinking | $u_r(r,\mu) = \left(-1 + \frac{3a}{2r} - \frac{a^3}{2r^3}\right) U\mu, \quad u_\theta(r,\mu) = \left(1 - \frac{3a}{4r} - \frac{a^3}{4r^3}\right) U\sqrt{1-\mu^2}$ |

| | **Fluid velocity field in lab frame** | **Far-field signature** |
|---|---|---|
| Sessile | same as above | Force monopole ($u \sim 1/r$) (Stokeslet) |
| Swimming | $u_r(r,\mu) = \frac{2a^3}{3r^3} B\mu,\ u_\theta(r,\mu) = \frac{a^3}{3r^3} B\sqrt{1-\mu^2}$ | Potential dipole ($u \sim 1/r^3$) |
| Sinking | $u_r(r,\mu) = \left(\frac{3a}{2r} - \frac{a^3}{2r^3}\right) U\mu,$ $u_\theta(r,\mu) = \left(-\frac{3a}{4r} - \frac{a^3}{4r^3}\right) U\sqrt{1-\mu^2}$ | Force monopole ($u \sim 1/r$) (Stokeslet) |

Table 2: Table 1 extension

| | Fluid pressure field | Forces on sphere |
|---|---|---|
| Sessile | $p(r,\mu) = p_\infty - \eta \frac{a}{r^2} B\mu$ | $\mathbf{F} = 4\pi\eta a B \mathbf{e}_z$ |
| Swimming | $p(r,\mu) = p_\infty - \eta \frac{a}{r^2} B\mu$ | $\mathbf{F} = 4\pi\eta a B \mathbf{e}_z, \quad \mathbf{D} = -6\pi\eta a U \mathbf{e}_z$ |
| Sinking | $p(r,\mu) = p_\infty + \eta \frac{3a}{2r^2} U\mu$ | $\mathbf{W} = -\frac{4}{3}\pi a^3 (\delta\rho) g \mathbf{e}_y, \quad \mathbf{D} = 6\pi\eta a U \mathbf{e}_y$ |

| | Hydrodynamic Power | Swimming Speed |
|---|---|---|
| Sessile | $\mathcal{P} = 8\pi a\eta B^2$ | $U = 0$ |
| Swimming | $\mathcal{P} = 8\pi a\eta \frac{2}{3} B^2$ | $U = \frac{2}{3} B$ |
| Sinking | $\mathcal{P} = 6\pi a\eta U^2$ | $U = \frac{2ga^2(\delta\rho)}{9\eta}$ |

Table 3: Expressions for Sh as a function of Pe for sessile and swimming ciliated sphere model, compared to a sinking sphere.

| | Large Pe limit | | Small Pe limit | |
|---|---|---|---|---|
| | Sherwood number | Reference | Sherwood number | Reference |
| Sessile | Sh $= \frac{2}{\sqrt{3\pi}} \mathrm{Pe}^{\frac{1}{2}}$ | present study | Sh $= 1 + \frac{43}{720} \mathrm{Pe}^2$ | present study |
| Swimming | Sh $= \frac{2}{\sqrt{3\pi}} \mathrm{Pe}^{\frac{1}{2}}$ | [14], [15] | $\mathrm{Sh} = 1 + \frac{1}{3} \mathrm{Pe}$ | [14], [15] |
| Sinking | Sh $= 0.55 \mathrm{Pe}^{\frac{1}{3}}$ | [17–19] | $\mathrm{Sh} = 1 + \frac{1}{3} \mathrm{Pe}$ | [17, 19] |

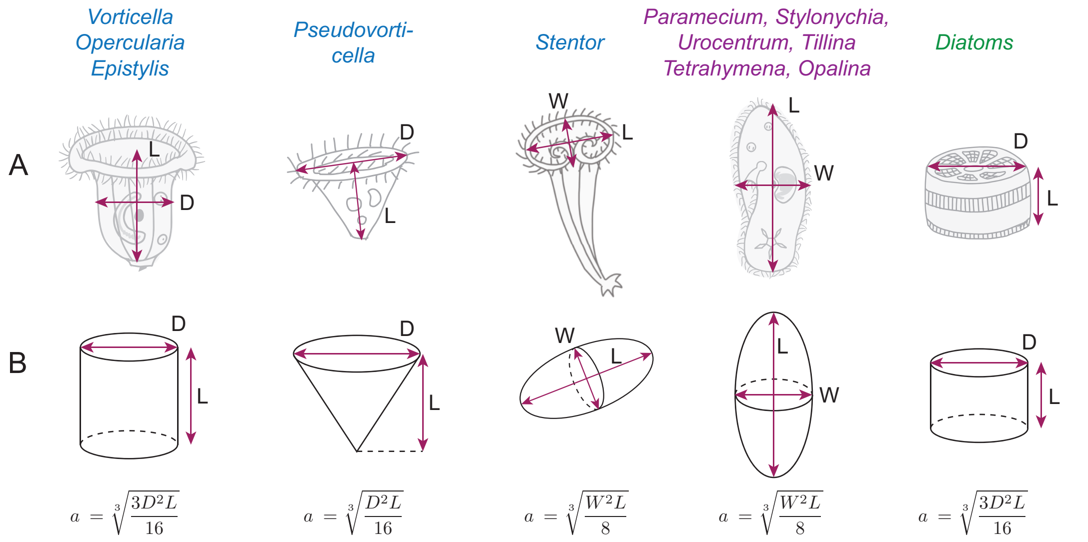


Figure 1: **Geometry of ciliates and diatoms**. (A) Representative morphologies of surveyed species of sessile ciliates (blue), swimming ciliates (purple), and sinking diatoms (green). (B) Simplified shapes of above organisms. The volumes of those shapes can be obtained as $V_{\text{cylinder}} = \pi D^2 L/4$, $V_{\text{cone}} = \pi D^2 L/12$, and $V_{\text{spheroid}} = \pi W^2 L/6$. Later on, we further simplify those shapes to a equal volume based sphere with radius $a$, calculated as formula shown in B for each geometry.

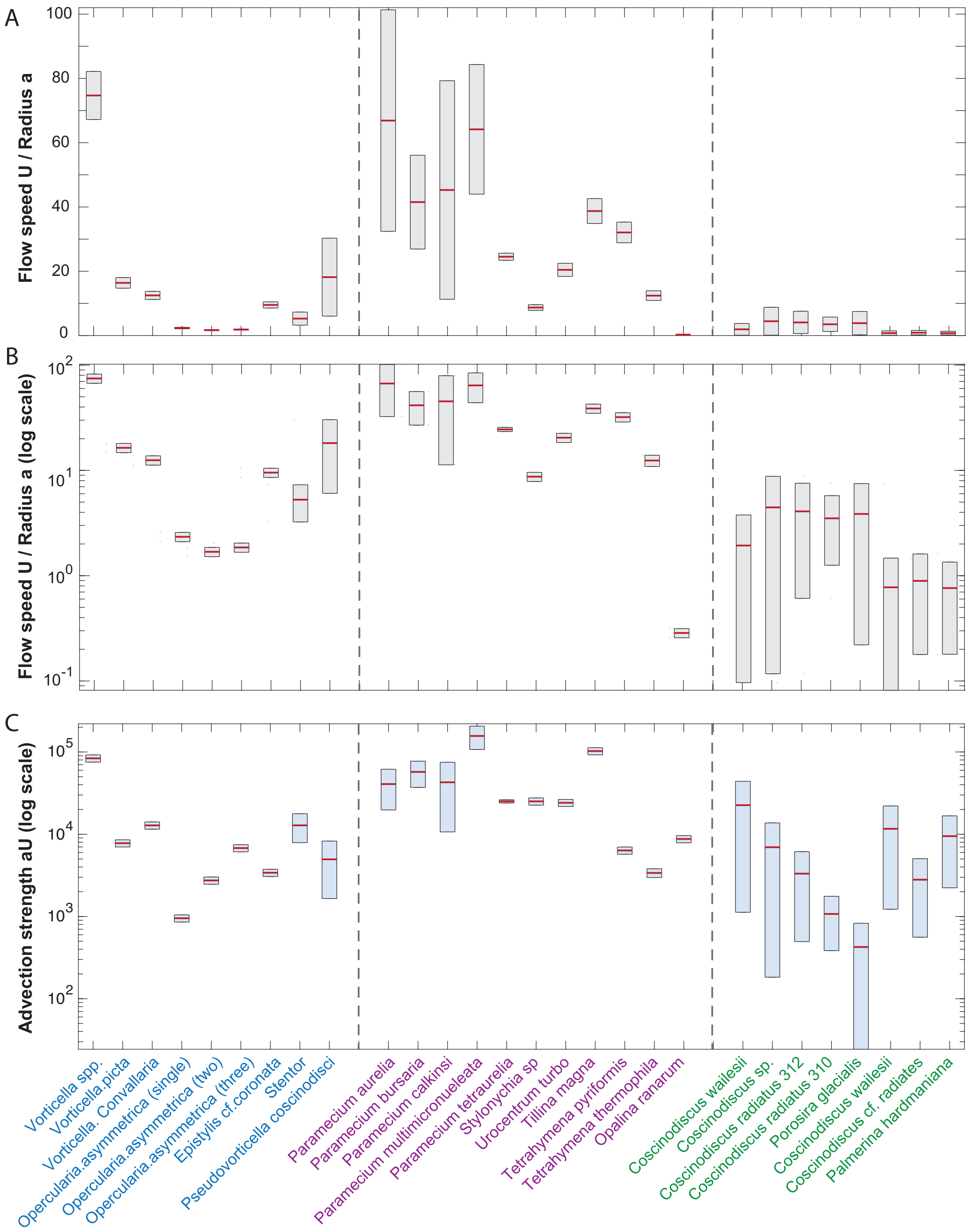


Figure 2: **Raw data of character flow speed and size** for sessile (blue) and swimming (purple) ciliates, and sinking diatoms (green). For motile organisms, the characteristic flow speed $U$ is the swimming speed, while for sessile organisms, the characteristic speed $U$ is the maximum flow speed reported near the organism. The characteristic length $a$ is based on the volume-equivalent spherical radius. (A) Ratio of Flow speed to length scale $U/a$ in regular scale. (B) Ratio of Flow speed to length scale $U/a$ in logarithmic scale. (C) Advection strength $aU$ in logarithmic scale.

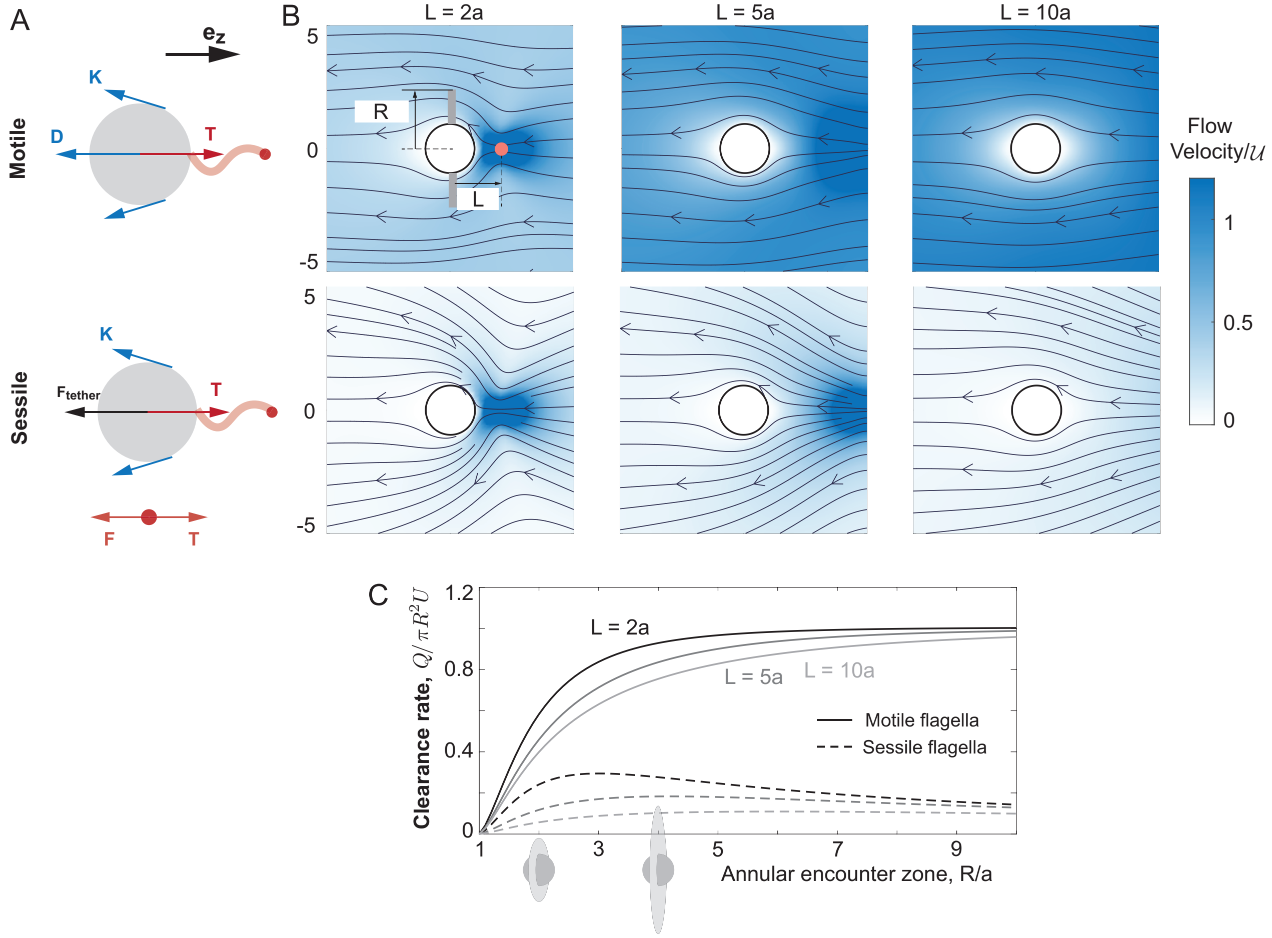


Figure 3: **Stokeslet model.** Flow field around sessile and swimming sphere. Solution in non-dimensional form for $a = 1$ and $B = F_{\text{cilia}}/(8\pi\mu a) = 1$.(A) Fluid around a motile and sessile sphere with same point force strength at distance $L = [2a, 5a, 10a]$ respectively. (B) Normalized clearance varies as annular encounter disk $R$ for above same three point force distances $L = [2a, 5a, 10a]$.
